# Mitochondrial Derived Vesicles retain membrane potential and contain a functional ATP synthase

**DOI:** 10.1101/2022.07.10.499450

**Authors:** Reut Hazan (Ben-Menachem), Dvora Lintzer, Tamar Ziv, Koyeli Das, Irit Rosenhek-Goldian, Ziv Porat, Hila Ben Ami Pilo, Sharon Karniely, Ann Saada, Neta Regev-Rudzki, Ophry Pines

## Abstract

Vesicular transport is a means of communication. While cells can communicate between each other via secretion of extracellular vesicles, less is known regarding organelle-to organelle communication, in particularly in the case of mitochondria. Mitochondria are responsible for the production of energy and for essential metabolic pathways in the cell, as well as fundamental processes such as apoptosis and aging. Here we show that functional mitochondria, isolated from *Saccharomyces cerevisiae* release vesicles, independent of the fission machinery. We were then able for the first time to isolate these Mitochondrial Derived Vesicles (MDVs) and found that they are relatively uniform in size, of about 100nm and carry selective protein cargo including enrichment of ATP synthase subunits. Remarkably, we further found that these MDVs harbor a functional ATP synthase complex. Moreover, we demonstrate that these vesicles have a membrane potential, produce ATP, and seem to fuse with naive mitochondria. Our findings reveal a possible delivery mechanism of ATP producing vesicles, which can potentially regenerate ATP deficient mitochondria and may participate in organelle to organelle communication.

## Introduction

Eukaryotic cells are defined by the existence of subcellular compartments and organelles, which allow partitioning of various biochemical pathways out of the cytosolic milieu into discrete organelles. In this sense, each cellular compartment has a specific protein composition that is vital for its function. Molecules of one protein can be located in several subcellular locations, a phenomenon termed dual targeting or dual localization. These identical or nearly identical forms of proteins, localized to different subcellular compartments are termed echoforms or echoproteins (to distinguish them from isoforms/isoproteins) (Kalderon and Pines, 2014; Morgenstern et al., 2017; Yogev and Pines, 2011). Dual targeting has been shown to be highly abundant and we now estimate that in yeast, more than one-third of the mitochondrial proteome is dual-targeted (Ben-Menachem et al., 2011; Kisslov et al., 2014). The vast majority of mitochondrial proteins are nuclear encoded, and are synthesized on cytosolic polysomes. A composite group of cytosolic and mitochondrial proteins take part in recognition, targeting and translocation of mitochondrial proteins from the cytosol to their destined organelle. Within mitochondria, proteins are sorted to one of their four sub-compartments: The outer membrane (OM), the intermembrane space (IMS), the inner membrane (IM) and the matrix. While there is thorough research on import of proteins into mitochondria, protein export from the organelle is perceived as nonexistent (Grevel et al., 2019; Neupert and Herrmann, 2007; Schneider, 2020).

Mitochondria occupy a substantial portion of the cytoplasmic volume of eukaryotic cells, and they have been essential for the evolution of eukaryotes. Mitochondria fulfill many crucial functions in the eukaryotic cell such as oxidative ATP production, providing building blocks for biosynthesis of macromolecules, β-oxidation of fatty acids, heme biosynthesis and formation of iron-sulfur (Fe-S) clusters (Reichert and Neupert, 2004). Apart from their normal physiological functions, mitochondria also play an important role in pathological phenomena in cells, including programmed cell death (apoptosis) (Jiang and Wang, 2004) and ageing (Balaban et al., 2005). All these reactions involve flow of substrates and products between the organelles within the cell, and direct inter-organellar contact is frequently required. Membrane contact sites (Scorrano et al., 2019) is a major mode of interaction between organelles and a means to maintain cellular homeostasis (Rizzuto et al., 1998; Zhang et al., 2005). Another possible mode of mitochondrial cross-talk was indicated by a report that mitochondrial cargo proteins are sorted into, at the time, uncharacterized mitochondrially-derived vesicles (Neuspiel et al., 2008). Mitochondrial derived vesicles (MDVs) can be generated either from the outer membrane of mitochondria, and may include both outer, inner membranes and matrix content (Neuspiel et al., 2008; Soubannier et al., 2012a; Soubannier et al., 2012b). These MDVs are defined by their diameter (between 70 and 150 nm), their independence of their formation on the mitochondrial fission GTPase Drp1 (Dnm1 in yeast) and by selective incorporation of protein cargo (Soubannier et al., 2012b). Of particular note is that MDVs formed *in vitro* are selectively enriched in oxidized cargo and are bound for late endosomes/lysosomes for degradation(McLelland et al., 2014; Neuspiel et al., 2008). In yeast and fungi vesicles termed ectosomes were reported in 2007 (Rodrigues et al., 2007), but since then little has been done to unravel the structure and function of these vesicles.

In the current research, we wished to gain understanding regarding the function of MDVs in protein localization and signaling, and to this end, we developed a method to isolate the secreted vesicles from yeast mitochondria. We then confirm that yeast mitochondria can generate vesicles which display selectivity in protein cargo including several ATPase subunits. Moreover, we describe for the first time the presence of an active ATPase machinery in these vesicles and their capability to generate ATP independently, and remarkably transfer this ability to other naïve mitochondria.

## Results

### Isolated and purified mitochondria from S. cerevisiae generate vesicles

In order to determine the function of MDVs in signaling, it was essential to purify large amounts of these vesicles and determine their cargo. To this end, we characterized and investigated the role of MDVs from *S. cerevisiae*. Initially, we isolated mitochondria from yeast cells using standard isolation methods (Daum et al., 1982), following a further purification step by percoll gradient, in order to separate mitochondria from contaminating organelles (e.g. ER) (Cunningham and Wickner, 1989; Trotter and Voelker, 1995). The purity of our isolated mitochondria was monitored with antibodies against mitochondrial [heat shock protein 60 (Hsp60)] and cytosolic [hexokinase 1 (Hxk1)] markers. As shown in figure 1A, the mitochondrial fraction is essentially free of cytosolic contamination, indicating the efficiency of our isolation method. Moreover, the quality of our isolated mitochondria was examined by an *in-vitro* import assay(Daum et al., 1982) of the TCA cycle protein, aconitase (Aco 1). In this experiment (Figure 1B), the labeled in vitro-transcribed Aco1 precursor, was incubated with our pure mitochondria, and the efficiency of the import process, was monitored by the existence of the mature form of the protein. As shown in figure 1B, the mature form of the protein was detected after 2 minutes, while after 15 minutes, almost 50 percent of the protein population was processed, indicating that the mitochondria are intact and have a membrane potential. Taken-together, these results indicate that our mitochondria are pure and active and suitable for MDVs isolation.

**Figure 1.**
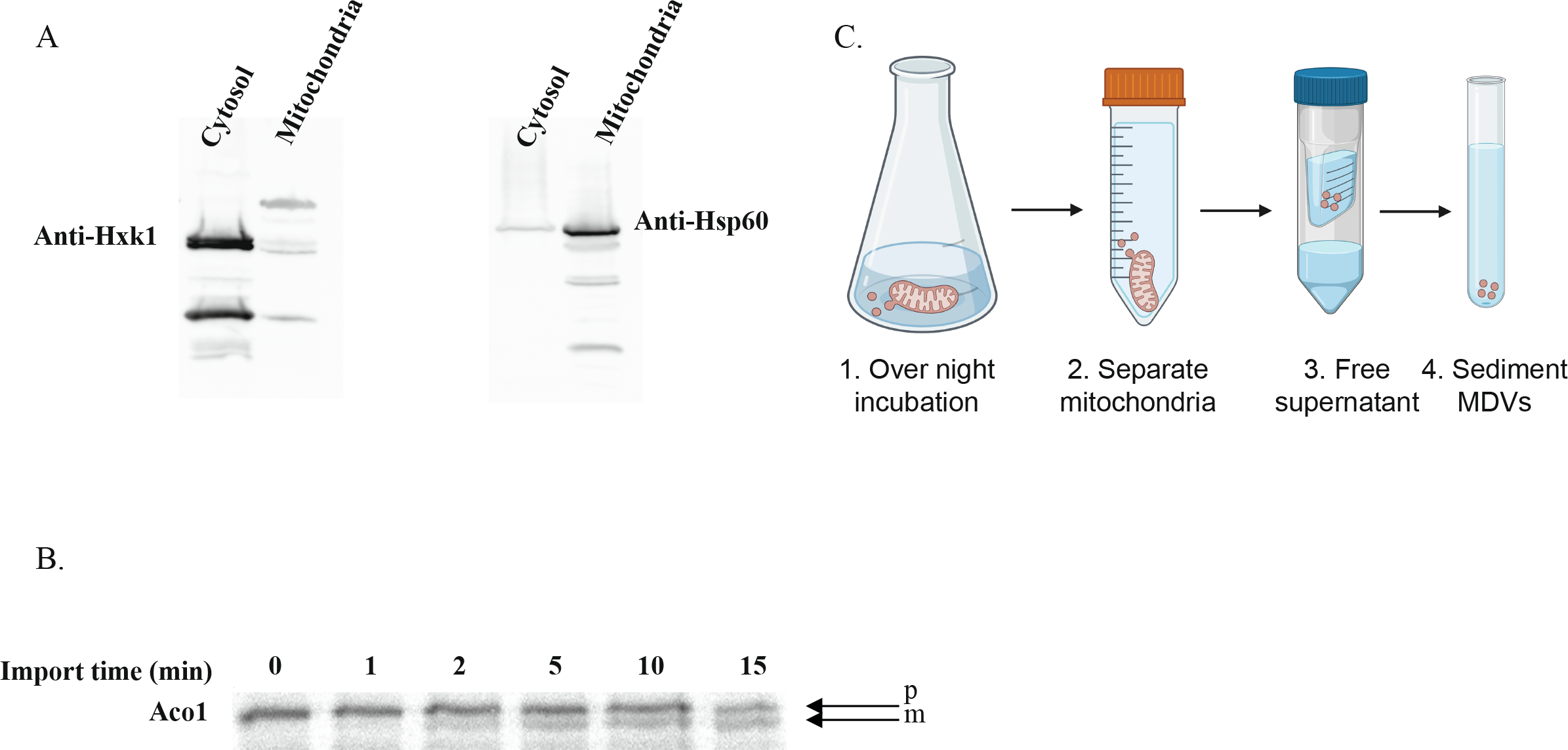
**A. Western blot analysis of purified mitochondria.** Mitochondria were isolated from yeast BY4741 wild type strain, and their purity was assessed by western blot analysis using Hsp60 and Hxk1 antisera as mitochondrial and cytosolic markers, respectively. **B. In-vitro import of Aco-1**. Aco1 mRNA was translated in rabbit reticulocyte lysate in the presence of [^35^S] methionine. The labeled protein products were added to purified mitochondria for the indicated times, in order to measure the mitochondrial import efficiency. Samples were analyzed by SDS-PAGE and autoradiography. Processing of the Precursor, p form to a Mature, m form, indicated a functional import machinery. (Vitrobot mark IV, FEI). The vitrified samples were examined at −177 °C using FEI Tecnai 12 G2 TWIN TEM operated at 120 kV and equipped with a Gatan model 626 cold stage. A 4K x 4K FEI Eagle CCD camera was used to record the images in low dose mode. **C. Representation of the vesicles isolation method**. Isolated and purified mitochondria were incubated in an isotonic buffer (1) and separated from the supernatant by centrifugation (2). The supernatant was freed from debris by viva-cell centrifugation (through vertical filtration membranes)(Sisquella et al., 2017) (3), which was then ultracentrifuged at high speed to sediment MDVs (4).

Previous work (Neuspiel et al., 2008) suggested that MDVs should be defined by at least three criteria;

I) relatively uniform diameter of 70 to 150 nm, II), independence of the mitochondrial fission protein Drp1, and III) selective incorporation of protein cargo. We proceeded to isolate yeast MDVs (Figure 1C) and examined our results according to these criteria. In order to determine whether mitochondrial derived vesicles are released from mitochondria, we used differential centrifugation to distinguish MDVs from intact mitochondria (Daum et al., 1982). Isolated and purified mitochondria were incubated in an isotonic buffer and separated from the supernatant by centrifugation. The supernatant was freed from debris by viva-cell centrifugation (through vertical filtration membranes)(Sisquella et al., 2017), which was then centrifuged at high speed (Figure 1C). Size and population analysis were determined by using four different strategies, as shown in figure 2. Nanosight particle analysis, revealed a particle diameter distribution between 80nm to 200nm with a peak at 105.8 +/− 2.2nm, which corresponds to the typical size of mitochondrial vesicles (Figure 2A) (Neuspiel et al., 2008). Biophysical analyses of mitochondrial derived vesicles by atomic force microscopy (AFM) and Transmission electron microscopy (TEM), verified that these vesicles are within a range of 50 to 200nm diameter (Figure 2B and 2C respectively). Moreover, Cryo-transmission electron microscopy (cryo-TEM) showed that these vesicles are composed of single or potentially double membranes (Figure 2D) as described previously(Neuspiel et al., 2008). Together, these results indicate that vesicles released by isolated mitochondria are of the expected size. To determine whether mitochondrial vesicle formation requires the mitochondrial fission GTPase Dnm1, we isolated mitochondria from KO cells of this protein and repeated vesicle isolation procedure. Size and population analysis were determined as described (Figure 2) and the results were similar to the *wild type* strain (Supplementary figure 1), indicating vesicle scission does not require this fission machinery.

**Figure 2.**
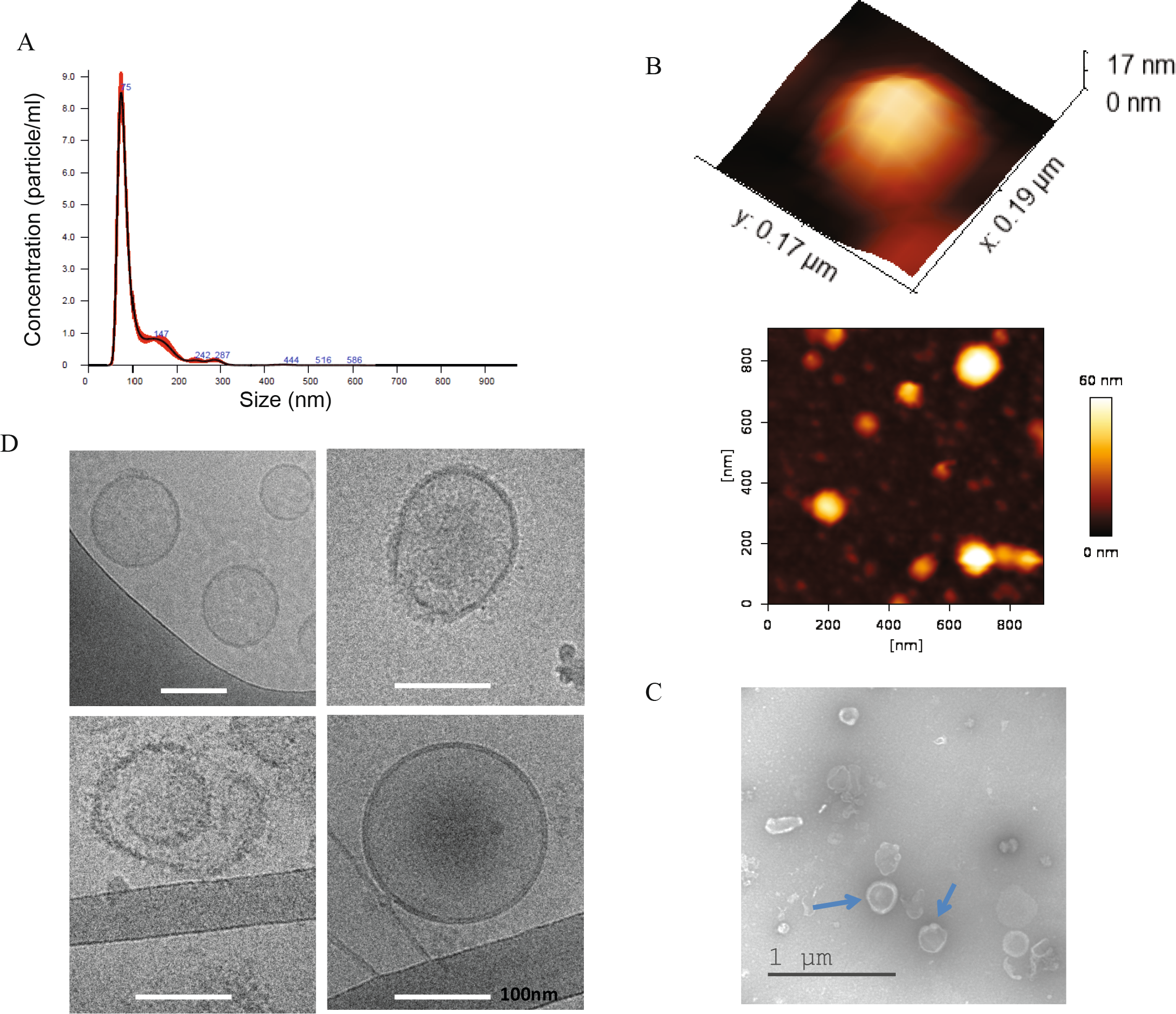
**A. Nanoparticle tracking analysis.** Vesicle size distribution and concentration was performed using Nanoparticle Tracking Analysis (NTA) (Malvern Instruments, Nanosight NS300). Sample size distributions were calibrated in a liquid suspension by the analysis of Brownian motion via light scattering. Nanosight provides single particle size and concentration measurements. **B. Atomic Force Microscopy**. Representative AFM image and a 3D AFM image of one representative vesicle (WT), adsorbed on a mica modified with Mg2+” and imaged under PBS. **C. TEM images of MDVs**. Samples were stained with 2.0% uranyl acetate or 2.0% phosphotungstic acid. 5 μl of vesicles was placed on Formvar/carbon coated copper 200 mesh grids (EMS), mixed with 5 μl of PTA for 10-20 sec, while excess stain was blotted off and grids were dried. Samples were examined with Jeol (Jem-1400 Plus) transmission electron microscope. **D. Cryo-TEM analysis of Mitochondrial derived vesicles**, which can be detected to contain single and potentially double membranes. Samples were prepared using a vitrification robot system. Scale bar– 100nm.

### MDVs show selectivity in protein cargo

It is well known that MDVs and extracellular vesicles (EVs) contain a variety of proteins and nucleic acid cargoes (Hernandez-Barranco et al., 2021; Neuspiel et al., 2008),(Abels and Breakefield, 2016; Gezsi et al., 2019; Ofir-Birin et al., 2017). In order to determine cargo selectivity of MDVs, isolated vesicles were purified by an OptiPrep (OP) gradient, an established method for EV purification (Regev-Rudzki et al., 2013), and fractions were loaded on acrylamide gels for validation of gradient efficiency. As shown in figure 3A, fractions 3 to 5 in which most MDVs were concentrated, were chosen for further analysis by LC/MS/MS. Comparison between the LC/MS/MS results of three independent repeats (Figure 3B Venn diagrams) led to identification of 667 mitochondrial proteins in the mitochondrial fractions, from which 521 were detected in all three repeats (Figure 3B, top right Venn diagram and Supplementary Table 1). The summed intensity of the mitochondrial proteins was more than 95% of the total intensity of the proteins. Our analysis revealed that out of these 667, only 207 mitochondrial proteins were identified in MDVs, from which 188 were detected in all three repeats (Figure 3B, top left and bottom Venn diagrams, Supplementary Table 2). The summed intensity of the mitochondrial proteins of the MDVs were more than 95% of the total intensity of the proteins.

**Figure 3.**
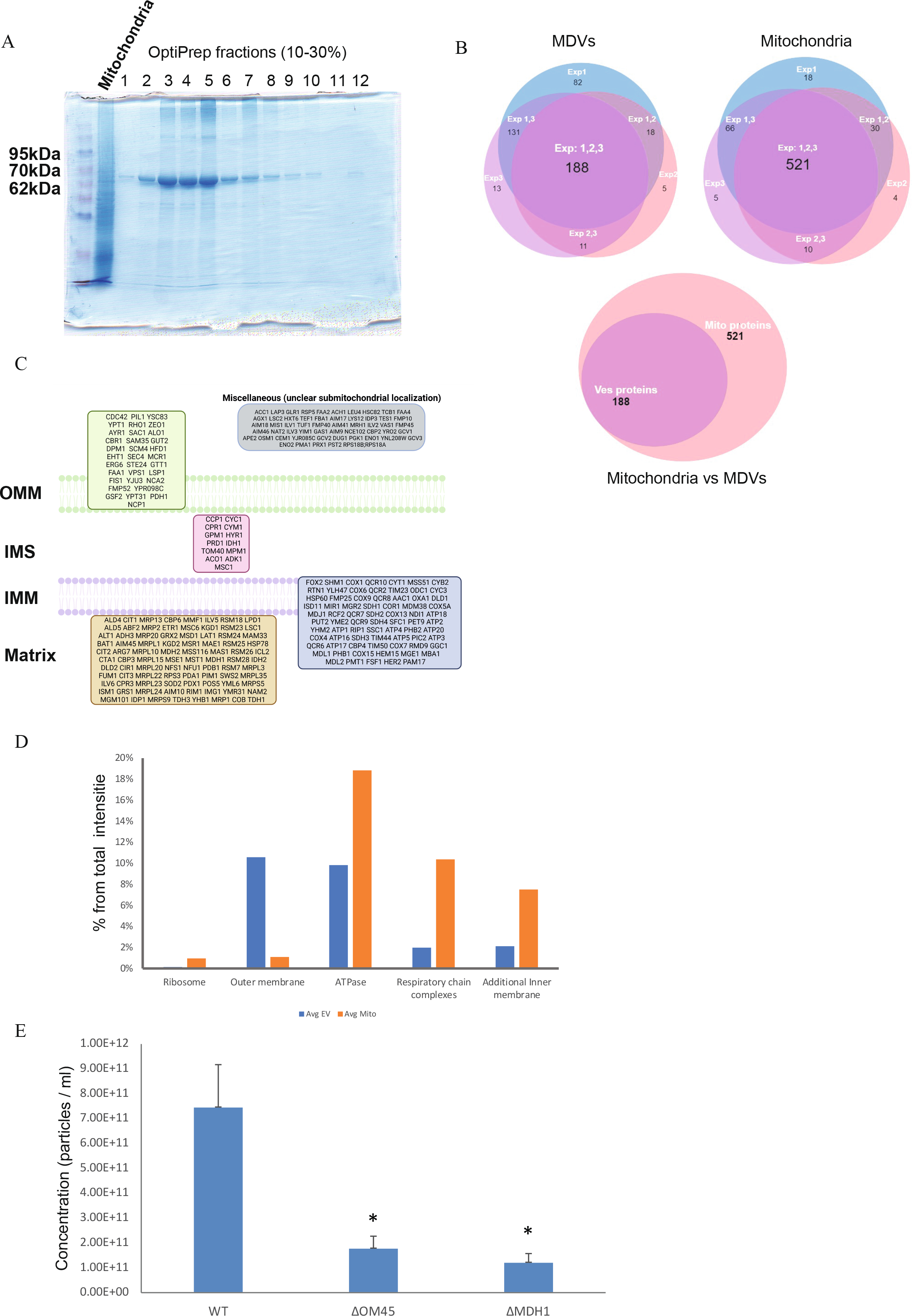
**A. OptiPrep Gradient Purification of MDVs.** Media components were fractionated by centrifugation (100,000g, 18hr, 4°C) through a continuous 10%–30% OptiPrep (Axis-Shield) gradient. Fractions (1 ml) were collected from the top of the gradient and run on acrylamide gel for further analysis. **B. Venn diagram of three different repeats of LC\MS\MS analysis**. 667 mitochondrial proteins were observed in mitochondrial fractions from three different repeats, and out of this group, 207 mitochondrial proteins were observed in MDVs. **C. Representation of sub-compartmental localization of MDVs proteome**. OMM– Outer Mitochondrial Membrane; IMM– Inner Mitochondrial Membrane; IMS– Inter Membrane Space. **D. There is a selectivity in protein cargo of MDVs versus Mitochondrial fractions**. Percentage of representative mitochondrial proteins from the total intensity of the identified proteins from three different repeats was calculated. Avg=average of three independent repeats. **EV= MDVs. Mito= mitochondria. E. Compromised mitochondria produce less MDVs**. MDVs isolated from Wild-type and KO strains (Δom45 and Δmdh1) were analyzed using Nanoparticle Tracking Analysis (NTA) (Malvern Instruments, Nanosight NS300). The results represent three independent repeats with p value< 0.05.

This group of 188 mitochondrial proteins includes inner and outer membrane proteins, as well as matrix soluble proteins, suggesting that vesicle proteins are acquired from internal mitochondrial sub-compartments (Figure 3C). As the protein profile of the vesicles were not similar to the profile of the Mitochondrial fractions, percentage of each protein from the total intensity of the identified proteins in the sample was calculated. Comparison between the vesicles and the mitochondrial fraction, revealed that proteins are unevenly distributed between those fractions. For example, the levels of ribosomal proteins in vesicles fractions were significantly lower than their levels in the mitochondrial fractions. Conversely, outer membrane proteins showed higher percentage from the total in the vesicles. (Figure 3D). Those differential protein profiles suggest that there is selectivity of mitochondrial proteins in MDVs. Specific examination shows proteins which are clearly identified in all three mitochondrial samples but absent from the MDVs samples (Supplementary Table 3). Conversely, there are a relatively few proteins clearly detected in vesicles (e.g., Icl1, according to number of peptides) but on the threshold of detection in the mitochondrial fraction. Those differential protein profiles suggest the presence of selected mitochondrial proteins in MDVs.

### MDV proteins affect vesicles formation and content

After characterizing the vesicles and examining their protein cargo, we attempted to approach the functional importance of such MDV proteins. Initially we asked whether mutations in genes encoding for proteins present in MDVs, would have an effect on the formation of these vesicles. In order to address this question, we isolated vesicles from two different KO strains; Mdh1 and Om45. Mdh1 is a mitochondrial matrix TCA enzyme while Om45 is an outer mitochondrial membrane protein. According to our mass-spec results, these proteins are found to be clearly detected in MDVs, and represent different locations of mitochondrial proteins that exist in MDVs (Supplementary Table 2). After validating by western blot analysis, the presence of Mdh1 in MDVs (Supplementary Figure 2), we compared MDV levels generated by these KO strains versus wild type strains, by Nanosight (NTA) particle analysis (Dekel et al., 2021; Gezsi et al., 2019). As shown in Figure 3E, vesicle concentration was lower in the two KO strains, as compared to the wild-type strain (pvalue<0.05), suggesting that these proteins may affect vesicle formation. Nevertheless, the effect of these KO mutations is most probably indirect and caused by effects on general mitochondrial function.

### Functional importance of MDVs

It is well known that vesicles can be transferred between cells and function in many different cellular activities, such as cell-cell communication, virulence, cargo and genetic material transfer etc, but less is known regarding the functions of MDVs (Andrade-Navarro et al., 2009; Picca et al., 2022; Ryan et al., 2020). To assess the functional importance of MDVs, we performed a GO annotation enrichment– analysis of biological processes using the STRING software, on the 188 mitochondrial proteins that were identified in our MDVs. As shown in figure 4A, a number of metabolic and biosynthetic pathways are represented in the vesicle proteome, while the ATP production related processes were one of the most significant. The mitochondrial F1Fo-ATP synthase produces the bulk of cellular ATP, and in yeast, it consists of 17 different structural subunits of dual genetic origin (Song et al., 2018). Thus, we decided to focus on this fascinating complex which comprises complex V of the mitochondrial respiratory chain. According to our mass-spec results, most subunits were detected in MDVs (supplementary table 2). To verify the presence of some of these subunits, we subjected yeast MDVs to western blot analysis, with anti-Atp2 antibody which is a crucial structural subunit of the mitochondrial ATP synthase. As shown in figure 4B, we detected Atp2 in all three main OptiPrep fractions of the secreted vesicles. Additionally, we examined the presence of Atp2 and Atp1 in vesicles by using immunogold labeling (Figure 4C). Isolated vesicles were first incubated with anti-Atp1 or anti-Atp2 antibodies, and then with 12nm gold goat anti rabbit antibodies. Samples were inspected by transition electron microscopy and as shown in figure 4C, our results confirmed that both Atp1 and Atp2 are present in vesicles, as compared to the appropriate controls (without the first antibody and anti-Hxk1 antibody, cytosolic protein, non-mitochondrial). Moreover, the average number of gold particles of Atp2 was significantly higher compared to the controls [pValue<0.05] (Supplementary Figure 3). Together, these results further support the presence of F1Fo-ATP synthase subunits in our vesicles.

**Figure 4.**
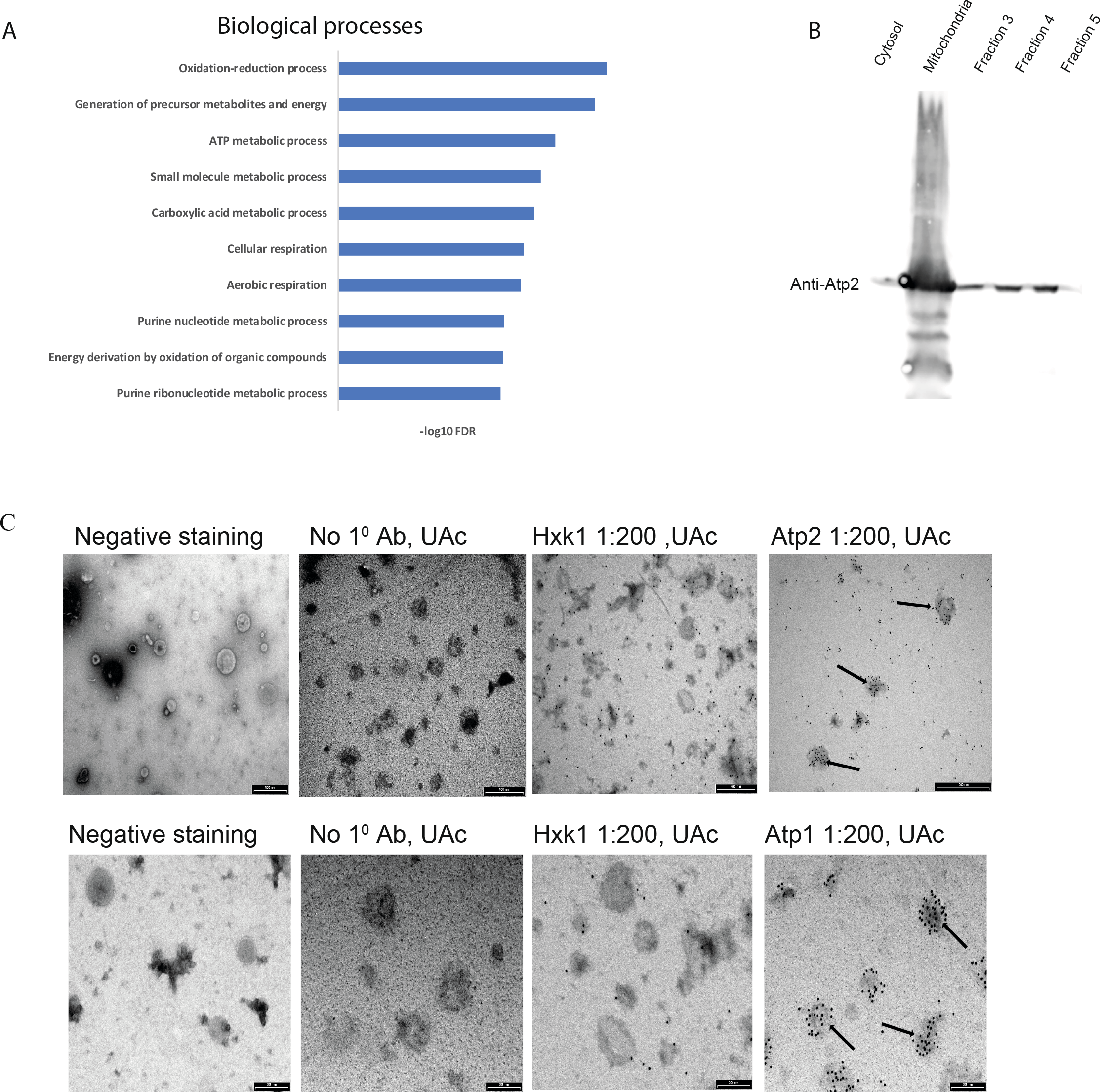
**A. GO annotation of biological processes of MDVs proteome.** A list of 188 MDVs proteins (based on three different repeats) were analyzed by the “STRING” tool. The results were filtered by strength>0.5 and FDR< 0.05. Here we represent the top 10 high significant values. **B. Western blot analysis of gradient purified MDVs**. Media components were fractionated by centrifugation (100,000g, 18hr, 4°C) through a continuous 10%–30% OptiPrep (Axis-Shield) gradient. Fractions (1 ml) were collected from the top of the gradient, ran on acrylamide gels and assessed by western blot analysis using Atp2 antisera. **C. Immunogold labeling of MDVs**. Isolated mitochondrial derived vesicles were incubated with specific primary antibodies (anti-Atp1 or anti-Atp2) and secondary antibody conjugated to 12nm gold (goat anti rabbit). Samples were viewed by Tecnai 12 TEM 100kV (Phillips, Eindhoven, the Netherlands). AUc– uranyl acetate. Scale bar– 500nm.

We hypothesized that if MDVs contain functional F1Fo-ATP synthase in their membranes, it is possible that they may be able to generate ATP. If this intriguing possibility is true, one may ask if vesicles would be capable to transfer this ability to generate ATP between healthy and damaged mitochondria, and rescue their phenotype. To address these issues, isolated MDVs from *wild-type* mitochondria were incubated with or without ADP in sorbitol buffer isotonic for mitochondria, and after washing and centrifugation steps, ATP production was determined by luminometry using luciferin-luciferase. As shown in figure 5A, during the 15 min incubation time in 30°C, a significant amount of ATP was produced compared to the control (no added ADP). Notably, a decreased amount of ATP was produced at 4°C compatible with an enzymatic reaction. Moreover, in the presence of the uncoupler CCCP or the ATP synthase (complex V) inhibitor oligomycin, the reaction was partially but significantly inhibited, indicating mitochondrial ATP production. Furthermore, insignificant amounts of ATP were detected in the second wash, suggesting that no MDVs remained in the reaction mix. Taken together these results indicate that yeast MDVs can produce ATP in an enzymatic reaction, which is dependent on F1Fo-ATP synthase and membrane potential. The existence of a membrane potential was also corroborated using image stream analysis, as shown in figure 5B. Atp2-GFP vesicles treated with TMRE (tetramethylrhodamine, ethyl ester) showed accumulation of this dye, and moreover, adding the uncoupler CCCP, reduces TMRE accumulation, indicating the existence of an electrochemical proton gradient. While setting up the ATP production experiments, we noted to our surprise, that the addition of TCA cycle substrates such as succinate, pyruvate or glycerol had no or insignificant effects on vesicle ATP production (Figure 5A). Moreover, MDVs from yeast devoid of malate dehydrogenase (a TCA cycle enzyme) were able to efficiently drive production of ATP (Figure 5A). Accordingly, we deduce that the membrane potential, *per se* was sufficient to drive ATP production.

**Figure 5.**
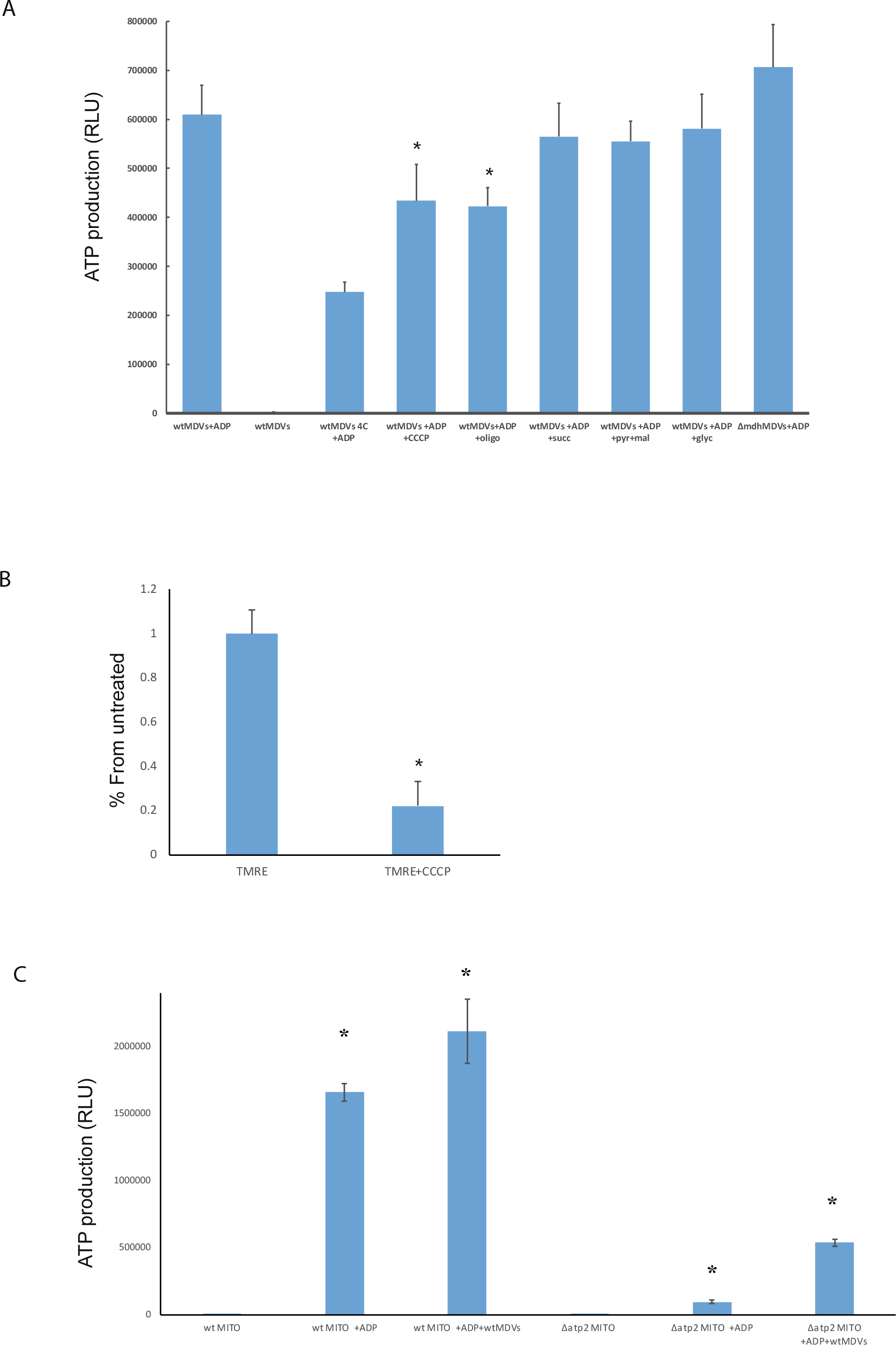
**A. MDVs produce ATP in an enzymatic reaction dependent on membrane potential.** Mitochondrial derived vesicles from wild type (WTves) or devoid of malate dehydrogenase (dMDHves), were incubated in the presence or absence of ADP, CCCP or oligomycin (oligo) for 15 minutes at 30°C or 4°C. Subsequently, ATP was measured by luciferin-luciferase luminometry. The results represent the average of three independent repeats. (Oligo– oligomycin, Succ-succinate, Pyr+Mal- pyruvate+malate, Glyc- glycerol). **B**. M**embrane potential of MDVs as visualized by Imaging Flow Cytometry**. Atp2-GFP MDVs stained with TMRE (tetramethylrhodamine, ethyl ester) were treated with CCCP and imaged by ImageStreamX mark II (Amnis, Part of Luminex, Au. TX). The two treatment show statistically significant difference (p-Value= 0.0099). **C. MDVs can be taken up by mitochondria and rejuvenate damaged mitochondria**. Isolated mitochondria from wild type (WT mito) and respiratory deficient mitochondrial (dATP2 mito) were preincubated in the presence or absence of mitochondrial derived vesicles from wild type (WTves), washed and incubated for 15 min at 30°C in the presence or absence of ADP. Subsequently, ATP formed was measured by luciferin-luciferase luminometry.

To assess whether wild type MDVs were able to compensate for impaired ATP production of F1Fo-ATP synthase deficient mitochondria, we preincubated wild type MDVs with wild type mitochondrial versus Atp2 respiratory deficient mitochondria (ΔAtp2), prior to the ATP production assay. As shown in figure 5C, ATP production by wild type mitochondria increased to some extent (30%). The major finding here was the effect of wild type MDVs on ΔAtp2 mitochondria which increased more than 5-fold (560%) (Figure 5C). Regretfully we could not perform the reciprocal experiments because we were not able to isolate (despite several attempts) MDVs from ΔAtp2 and we assume that most probably an intact F1Fo-ATP synthase may influence MDV production. Taken together, these results suggest that MDVs may rejuvenate damaged mitochondria by sending MDVs between healthy and damaged mitochondria.

To corroborate some of the above results in another system, we were able to isolate MDVs from isolated mitochondria derived from HEK293 cells and analyzed their size by Nanosight particle analysis. As shown in supplemented figure 4A, this analysis revealed a typical size of mitochondrial vesicles (Neuspiel et al., 2008). Moreover, biophysical analyses of mitochondrial derived vesicles Transmission electron microscopy (TEM), verified that these vesicles are within a range of 50 to 200nm diameter (supplemented figure 4C). Next, we measured ATP production in HEK293 derived MDVs and in accord we observed ATP production (supplemented figure 4C), which was also significantly decreased in the presence of CCCP or oligomycin (supplemented figure 4C).

### Discussion

One of the challenges when dealing with the upcoming field of Mitochondrial Derived Vesicles (MDVs), is not only to define their composition and specific structure, but as we show here, demonstrate an activity and function associated with the MDVs. This gives us confidence against the arguments that in some cases MDVs are simply ruptured degradation products of mitochondria.

Initially we characterized the proposed and highly purified MDVs by a number of approaches (NTA, Nanoparticle Tracking Analysis; TEM, transmission electron microscopy; AFM, Atomic Force Microscopy [3D]; Cryo-TEM analysis). These MDVs, indeed resemble classical mitochondrial secreted vesicles and have a relatively uniform size of about 100nm with single or double membranes. Another important point worth discussing, is that less than half of the identified mitochondrial proteome were detected in the MDVs. Specific examination of this list of proteins shows that there are 72 proteins were nicely identified in all 3 experiments but are absent from the MDVs. These results suggest that there appears to be a selection of mitochondrial proteins as vesicle cargo.

Of the number of ATP production related processes, represented in the vesicle proteome, the mitochondrial F1Fo-ATP synthase was clearly outstanding. According to our results, most subunits were detected by mass-spec of MDVs and we confirmed the presence of Atp2 by western blot analysis. Additionally, we detected the presence of Atp2 and Atp1 in vesicles by immunogold labeling.

The major discoveries in this study are that MDVs have a membrane potential and a functional F1Fo ATP synthase complex with the capability to produce ATP. Like mitochondrial complex V, MDVs ATP production is sensitive to membrane potential dissipaters and ATP Synthase inhibitors. Thus, the MDVs acquire a major activity and function of mitochondria and, moreover may possibly transfer this ability from one organelle to the other. This process may allow rejuvenation of damaged mitochondria. In addition, ATP producing vesicles, may be directed to subcellular locations that require ATP such as the yeast bud or along axons in neurons without the need to translocate whole mitochondria.

Another line of thought involves protein targeting or as alluded to earlier, MDVs may be a pathway to export proteins from mitochondria for which such a pathway has not been identified by any other mechanism. In fact, our group has been interested in protein dual targeting or dual localization of mitochondrial proteins and MDVs may be a mechanism by which this can achieved. In this regard, there are reports identifying ATP synthase in the plasma membrane which could be achieved via an MDV driven pathway (Cavelier et al., 2012).

We know (According to our data) that mutations in some mitochondrial protein encoding genes (e.g. MDH1, OM45) can affect the amount of vesicles produced by mitochondria but assume that this is an indirect effect resulting from general mitochondrial wellbeing. We have no idea at present which proteins are mechanistically involved in MDV formation. Furthermore, in this study we have focused on protein biochemistry, cell biology and function of MDVs, however, there are many other aspects of MDV biology which will addressed by our group in the future.

## Materials and methods

### Yeast strains

*Saccharomyces cerevisiae* strains used were: BY4741 (Mat a, his31, leu20, met150 and ura30), ΔDnm1 (BY4741; MATa; his3Δ1; leu2Δ0; met15Δ0; ura3Δ0; LL001w∷kanMX4), ΔPor1 (BY4741; MATa; his3Δ1; leu2Δ0; met15Δ0; ura3Δ0; YNL055c∷kanMX4), ΔOm45 (BY4741; MATa; his3Δ1; leu2Δ0; met15Δ0; ura3Δ0; YIL136w∷kanMX4), ΔMdh1 (BY4741; MATa; his3Δ1; leu2Δ0; met15Δ0; ura3Δ0; YKL085w∷kanMX4), ΔAtp1 (BY4741; MATa; his3Δ1; leu2Δ0; met15Δ0; ura3Δ0; YBL099w∷kanMX4), ΔAtp2 (BY4741; MATa; his3Δ1; leu2Δ0; met15Δ0; ura3Δ0; YJR121w∷kanMX4), Atp2-GFP (BY4741; MATa; his3Δ1; leu2Δ0; met15Δ0; ura3Δ0; ATP2-GFP∷HIS).

### Growth Media

In order to isolate mitochondria, all strains were grown overnight at 30 °C in synthetic depleted (SG) medium containing 0.67% (w/v) yeast nitrogen base, 2% (w/v) galactose medium, supplemented with the appropriate amino acids (50 μg/ml). The next day the cells were washed and transferred to lactate medium containing 0.67% (w/v) yeast nitrogen base, 2% (v/v) lactate, 0.05% (w/v) glucose, supplemented with the appropriate amino acids (50 μg/ml) and adjusted to pH 5.5 with KOH.

### Western blot assay

Proteins were separated on 10% SDS-PAGE and transferred to nitrocellulose membranes. The primary antibodies used for detection were generously provided by Alexander Tzagoloff (Columbia University, USA) [Anti-Atp1 and anti Atp2], Doron Rappaport (University of Tubingen, Germany) [Anti Mdh1 and anti Por1]. Anti-Hxk1 was obtained commercially from ROCKLAND (catalog number 100-4159) and Anti-Hsp60 was generated in our lab. Secondary antibodies used are anti-Rabbit IgG-HPR (111-035-003, Jackson, dilution 1:10000). Western blot analyses were repeated at least three times. Image collection was performed using the ThermoScientific MyECL Imager V. 2.2.0.1250 and Amersham Imager 680 (GE Healthcare Life Sciences).

### In Vitro Translation and Import into Mitochondria

Transcription of pGEM plasmids in vitro was carried out according to the manufacturer’s instructions (Promega) using SP6 polymerase. Isolated mitochondria were prepared from yeast as described **(Daum et al., 1982)**. Protein synthesis was performed in the presence of [^35^S]methionine in rabbit reticulocyte lysate (Promega). Coupled translation/translocation reactions of 20 μl contained 16.5 μl of reticulocyte lysate, 0.5 μl of amino acid mix minus methionine, 0.5 μl of ribonuclease inhibitor, 1 μl of transcription reaction, and 10 μCi of [^35^S]methionine. In all experiments, translation was allowed to proceed for 2 min (unless otherwise indicated) before the addition of mitochondria (10–20 μg) and the reaction further incubated at 30 °C for 60 min. Following translation, all samples were diluted into SHKCl buffer (600 mm sorbitol, 50 mm Hepes, pH 7.2, 80 mm KCl). Selected samples were treated with 70 μg/ml proteinase K for 15 min on ice. Centrifugation at 9000 × g for 10 min at 4 °C yielded a supernatant fraction, and a mitochondrial pellet, which was washed once before being resuspended in SHKCl buffer. Samples were denatured by boiling in loading buffer containing 1% SDS, 4% β-mercaptoethanol, 1% dithiothreitol and analyzed on 10% SDS-polyacrylamide gel electrophoresis followed by visualization with the image-analyzing system BAS2000 (Fuji Corp.).

### Purification of Mitochondrial Derived Vesicles

Briefly, yeast mitochondria were purified and isolated as described previously (Daum et al., 1982). Next, isolated mitochondria were resuspended in SH buffer (0.6M sorbitol, 20mM hepes pH 7.4) for overnight in 30°C. Hek 293 Cell mitochondria were purified from eighteen confluent T75 flasks of HEK293T (ATCC) cells grown in DMEM with 10% fetal bovine serum supplemented with glutamate and penicillin streptomycin in 5% CO_2_ at 37°C. The cells detached by cell scraper and washed twice in phosphate buffered saline by centrifugation at 1800 x g at room temperature. The resulting cell pellet, was suspended in 2ml ice-cold buffer A (320 mM sucrose, 5 mM Tris, 2 mm EGTA, pH 7.4) and homogenized on ice by 300 strokes at 400rpm with a potter-elvehjem glass-teflon homogenizer. Nuclei were removed by 3 min centrifugation at 2000 x g, 4°C and mitochondria were pelleted from the supernatant by 10 min centrifugation at 12000 x g. After an additional wash, the mitochondria were suspended in 10 ml buffer A containing 2mg/ml fatty acid-free bovine serum albumin and incubated overnight in 5% CO_2_ at 37°C. Subsequently mitochondrial pellets were removed by centrifugation at 18000 x g by two repetitive centrifugations. The supernatant was concentrated using a Vivaflow 100,000 MWCO PES (Sartorious Stedium) and centrifuged at 150,000 × g, 18 h to pellet MDVs. Aliquot were made and MDVs were stored in −20°C until further use.

### OptiPrep Gradient Purification of MDVs

The initial vesicle pellet was resuspended in 2 ml SH buffer and further purified on a 10-ml continuous 10–30% OptiPrep gradient made up in NTE buffer (137mM NaCl, 1mM EDTA and 10mM Tris, pH 7.4), 150,000 g, 18 h, 4 °C, in a SW41 rotor (Beckman Coulter, Fullerton, CA, USA) (Dekel et al., 2021). Fractions (1 ml) were collected from the top for further analysis.

### Nanosight particle analysis (NTA)

MDVs size distribution and concentration were performed using Nanoparticle Tracking Analysis (NTA) (Malvern Instruments, Nanosight NS300). Sample size distributions were calibrated in a liquid suspension by the analysis of Brownian motion via light scattering. Nanosight provides single particle size and concentration measurements.

### Atomic Force Microscopy

Freshly cleaved mica surface was incubated with 10mM MgCl_2_ solution for 2 minutes, then rinsed with 200μl PBS. 50μl of MDVs solution was placed on the Mg modified mica for 10 min. Additional 50μl PBS were added to the sample prior to scanning. Images were captured using the JPK Nanowizard III AFM microscope (Berlin, Germany) in QI mode using qp-BioAC- Cl- CB-2 or CB-1 probes, spring constants≈0.1-0.2 N/m (Nanosensors). Image analysis was performed using JPK-SPM data processing software (Nečas and Klapetek, 2012).

### Transition Electron Microscopy

Samples were fixed with Karnovsky’s fixative followed by staining with 2.0% of uranyl acetate. Briefly, 5 μl of MDVs was placed on Formvar/carbon coated copper 200 mesh grids (EMS) and mixed with 5 μl of uranyl acetate for 10-20 sec. Excess stain was blotted off and grids were dried. Samples were examined with Jeol® JEM-1400 Plus TEM (Jeol®, Tokyo, Japan), equipped with ORIUS SC600 CCD camera (Gatan®, Abingdon, United Kingdom), and Gatan Microscopy Suite program (DigitalMicrograph, Gatan®, UK).

### Negative staining of MDVs

Vesicles were isolated from mitochondria and re-suspended in SH buffer (0.6M sorbitol, 20mM hepes pH 7.4). The vesicles were adsorbed to Carbon-Formvar coated copper grids. Grids were stained with 1% (w/v) uranyl acetate and air-dried. Samples were viewed with Tecnai 12 TEM 100kV (Phillips, Eindhoven, the Netherlands) equipped with MegaView II CCD camera and Analysis**®** version 3.0 software (Soft Imaging System GmbH, Münstar, Germany).

### Immuno labeling of MDVs

Isolated vesicles were adsorbed to Carbon coated nickel grids. From this point all steps were performed in drops of different reagents on parafilm within a wet chamber. Grids were washed 5 times in a Tris-buffered saline (TBS) containing 20mM Tris-base, 0.9% NaCl, 0.5% BSA, 0.5% Tween-20 and 0.13% NaN3, pH 8.2, and then incubated 30 minutes in a blocking solution containing 5% normal goat serum (NGS) in TBS. Grids were then exposed to antibodies for 16 hours at 4°C. The antibodies were diluted 1:200 with 1% NGS in TBS. Girds were then washed 5 times with TBS. For immunogold labeling grids were incubated with 12nm colloidal gold goat anti rabbit (Jackson ImmunoResearch Laboratories) for 1.5 hours followed by 5 washes in TBS and two washes in H_2_O and then by a 2 minutes-fixation of 2% glutaraldehyde.

Contrasting was achieved with saturated aqueous uranyl acetate and lead citrate before air-drying. Samples were viewed as described for negative staining.

### Proteolysis and mass spectrometry analysis

The proteins in the gel were reduced with 3mM DTT in 100mM ammonium bicarbonate [ABC] (60°C for 30 min), modified with 10mM Iodoacetamide in 100mM ABC (in the dark, room temperature for 30 min) and digested in 10% acetonitrile and 10mM ammonium bicarbonate with modified trypsin (Promega) at a 1:10 enzyme-to-substrate ratio, overnight at 37°C. The resulted peptides were desalted using C18 stage tips Uniprot and were analyzed by LC-MS-MS. The peptides were resolved by reverse-phase chromatography on 0.075 × 180-mm fused silica capillaries (J&W) packed with Reprosil reversed phase material (Dr Maisch GmbH, Germany). They were eluted with linear 60 minutes gradient of 5 to 28% 15 minutes gradient of 28 to 95% and 15 minutes at 95% acetonitrile with 0.1% formic acid in water at flow rates of 0.15 μl/min. Mass spectrometry was performed by Q Exactive plus mass spectrometer (Thermo) in a positive mode(m/z 300–1800, resolution 70,000 for MS1 and 17500 for MS2) using repetitively full MS scan followed by collision induces dissociation (HCD, at 25 normalized collision energy) of the 10 most dominant ions (>1 charges) selected from the first MS scan. The AGC settings were 3×106 for the full MS and 1×105 for the MS/MS scans. A dynamic exclusion list was enabled with exclusion duration of 20 s. The mass spectrometry data was analyzed using the MaxQuant software 1.5.2.8 (1) for peak picking and identification using the Andromeda search engine, vs. the Saccharomyces cerevisiae section of the uniport database. Oxidation on Methionine and acetylation on the N-terminus were accepted as variable modifications and carbamidomethyl on Cysteine was accepted as static modifications. Minimal peptide length was set to six amino acids and a maximum of 3miscleavages was allowed. Peptide- and protein-level false discovery rates (FDRs) were filtered to 1% using the target-decoy strategy. Protein table were filtered to eliminate the identifications from the reverse database and from common contaminants. The data was quantified by label free analysis using the same software, based on extracted ion currents (XICs) of peptides enabling quantitation from each LC/MS run for each peptide identified in any of experiments. Annotation and statistical analysis were done using the Perseus (2) and the String software (https://string-db.org/). The mass spectrometry proteomics data have been deposited to the ProteomeXchange Consortium via the PRIDE [1] partner repository with the dataset identifier PXD035192.

**Username**, **Password**: jkrWcDvT.

### Filtering and normalization of LC/MS/MS results

In order to establish a list of identified mitochondrial proteins on the project, LC-MS/MS results from three different repeats were filtered by comparing the results to three sources of yeast mitochondrial annotation: SGD, Uniprot and Go annotation analysis of cellular compartment by using STRING gene ontology tools. Following this filtering, we keep analyzing only the mitochondrial proteins. We calculate the intensity relative percentage of each protein, from the total intensity of all mitochondrial proteins.

### ImageStream analysis

The ImageStreamX system is an advanced imaging flow cytometer, combining features of fluorescent microscopy and flow cytometry. Cells in suspension pass through the instrument in a single file where transmitted light, scattered light and emitted fluorescence are collected, at a rate of up to 5000 cells s−1. This is accompanied by a dedicated image analysis software (IDEAS), which allows advanced quantification of intensity, location, morphology, population statistics, and more, within tens of thousands cells per sample. It allows analysis of rare subpopulations in highly heterogenous samples and gives rise to novel applications that were difficult to achieve by either conventional flow cytometry or microscopy (Zuba-Surma et al., 2007).

### Imaging Flow Cytometry

Vesicles were stained as described above and imaged by ImageStream mark II (Amnis, Part of Luminex, Au. TX). At least 3*104 events were collected from each sample. Lasers used were 488nm (400mw), 561nm (200mw) and 785nm (5mw). Channels acquired were 2 (GFP), 4 (TMRE), 6 (Side scatter), and 1+9 for the bright-field image. Inter-channel spillover was compensated by acquiring single stained controls. Images were analyzed using IDEAS 6.3 software (Amnis, Part of Luminex, Au. TX). Debris and aggregates were removed by plotting the bright field area (in square microns) vs. the side scatter intensity (The light scattered 90 degrees of the sample). Only objects with low bright field and low side scatter were taken for the analysis. Positive events for GFP and TMRE were gated by plotting the Max Pixel values (the largest value of the background-subtracted pixels contained in the input mask) of the GFP and TMRE staining. The gate was set according to the single stained controls.

### ATP production and complementation

The ATP production assay was adapted to Yeast and HEK293 cell MDVs based on our previously described work for mitochondria (Shufaro et al., 2012). Briefly 10^11^ MDVs/ml (stored at −20C and thawed immediately before the assay) were incubated for 15 min at 30°C for yeast MDVs or 37 °C for cell MDVs in assay buffer (5 mM K_2_HPO_4_, 10 mM, 100 mM KCl, 5 mM MgCl_2_, 0.005 mM EDTA, 75 mM mannitol, 25 mM sucrose, and 0.6 mg/mg fatty acid-free bovine serum albumin, at pH 7.4) in the presence or absence of 0.6 mM ADP, 40mM pyruvate, 20mM malate, 11mM succinate, 4μM cyanide 3-chlorophenylhydrazone(CCCP), 1.5 μM oligomycin (Sigma-Aldrich-Merck, Darmstadt, Germany). Subsequently, relative ATP levels were determined by luciferin-luciferase ATPlite luminescence assay system, (Perkin Elmer, Akron, Ohio, USA) according to the manufacturer’s instructions. Luminescence measurements were performed with a Synergy HT microplate reader (Bio-Tek instruments, Vinooski, VT, USA). For complementation assays mitochondria 10μg/μl) were pre-incubated with MDVs for 30 minutes in assay buffer at 30°C, then washed twice in large volumes of SH buffer (0.6M sorbitol, 20Mm Hepes pH 7.4) following centrifugation at 18,000g for 20 minutes at 4°C. The mitochondria were subsequently resuspended in the original volume and incubated in the presence or absence of ADP for 15 min before assaying ATP.

## Acknowledgements

We thank Alexander Tzagoloff (Columbia University, USA) for generously providing antisera and Yael Freidman (Hebrew University, Israel) for performing Electron Microscopy Images. The research of NR-R is supported by the Benoziyo Endowment Fund for the Advancement of Science, the Jeanne and Joseph Nissim Foundation for Life Sciences Research and the Samuel M. Soref and Helene K. Soref Foundation. NR-R is the incumbent of the Enid Barden and Aaron J. Jade President’s Development Chair for New Scientists in Memory of Cantor John Y. Jade. NRR is grateful for the support from the European Research Council (ERC) under the European Union’s Horizon 2020 research and innovation program (grant agreement No. 757743), the Minerva Program support (grant number 714142), and the Israel Science Foundation (ISF) (Grant Application no. 570/21).

---

**Supplementary figure 1. Characterization of MDVs derived from Dnm1 KO mitochondria. A. Nanoparticle tracking analysis.** Vesicle size distribution and concentration was performed using Nanoparticle Tracking Analysis (NTA) (Malvern Instruments, Nanosight NS300). Sample size distributions were calibrated in a liquid suspension by the analysis of Brownian motion via light scattering. Nanosight provides single particle size and concentration measurements. **B. Atomic Force Microscopy**. Representative AFM image and a 3D AFM image of one representative vesicle (WT), adsorbed on a mica modified with Mg2+” and imaged under PBS. **D**. Cryo-TEM analysis **of Mitochondrial derived vesicles**, which can be detected to contain single and potentially double membranes. Samples were prepared using a vitrification robot system. Scale bar- 100nm. **C. TEM images of MDVs**. Samples were stained with 2.0% uranyl acetate or 2.0% phosphotungstic acid. 5 μl of vesicles was placed on Formvar/carbon coated copper 200 mesh grids (EMS), mixed with 5 μl of PTA for 10-20 sec, while excess stain was blotted off and grids were dried. Samples were examined with Jeol (Jem-1400 Plus) transmission electron microscope.

**Supplementary figure 2. Western blot analysis of isolated MDVs.** MDvs were purified from wild type or KO isolated mitochondria as mentioned and the presence of Mdh1p and Por1p was assessed by western blot analysis using Mdh1 and Por1 antisera.

**Supplementary figure 3. The average number of gold particles of Atp2 is significantly higher compared to the controls.** The number of gold particles in 10 fields was messured using ImageJ software (ImageJ (nih.gov)).

**Supplementary figure 4. A. Nanoparticle tracking analysis of HEK293-derived MDVs.** Vesicles derived from HEK293 isolated mitochondria were subjected to size distribution and concentration using Nanoparticle Tracking Analysis (NTA). **B. TEM images of HEK293-derived MDVs**. Samples were stained with 2.0% uranyl acetate or 2.0% phosphotungstic acid. 5 μl of vesicles was placed on Formvar/carbon coated copper 200 mesh grids (EMS), mixed with 5 μl of PTA for 10-20 sec, while excess stain was blotted off and grids were dried. Samples were examined with Jeol (Jem- 1400 Plus) transmission electron microscope.

**C. HEK293-derived MDVs, produce ATP in an enzymatic reaction dependent on membrane potential**. Mitochondrial derived vesicles from HEK293-isolated mitochondria, were incubated in the presence or absence of ADP, CCCP or oligomycin (oligo) for 15 minutes at 37°C. Subsequently, ATP was measured by luciferin-luciferase luminometry. The results represent the average of two different biological repeats.

**Supplementary Table 1. Proteins of the mitochondrial fraction.** Proteins from three independent biological repeats were identified and quantified by LC-MSMS followed by data analysis using the MaxQuant software. The mitochondrial protein table contains information on the identified proteins that were annotated as mitochondrial and were identified and quantified in at least 2 repeats.

**Supplementary Table 2. Proteins of the mitochondrial vesicles fractions.** The vesicle protein table, a subset of Table 1, contains proteins that were identified and quantified in all three biological repeats

**Supplementary Table 3. Exclusive mitochondrial proteins**

Mitochondrial proteins with quantified in all three mitochondrial fraction and in none of the MDVs fractions.

