## Supplementary figures and images for "Mitochondrial Derived Vesicles retain membrane potential and contain a functional ATP synthase"

### supplemented figure 1

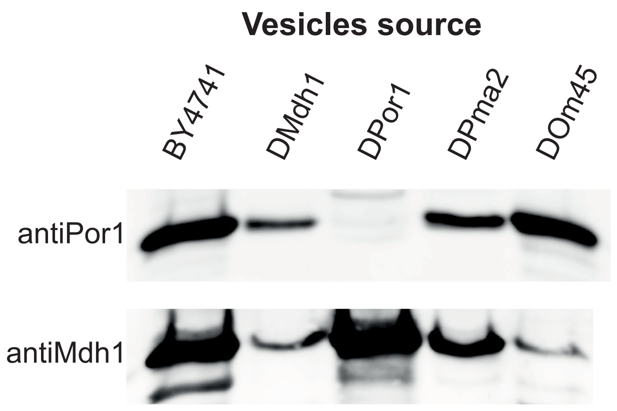

### supplemented figure 2

A.

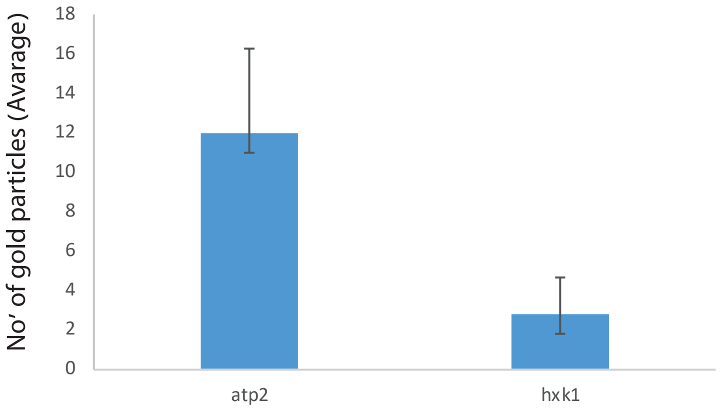

### supplemented figure 3

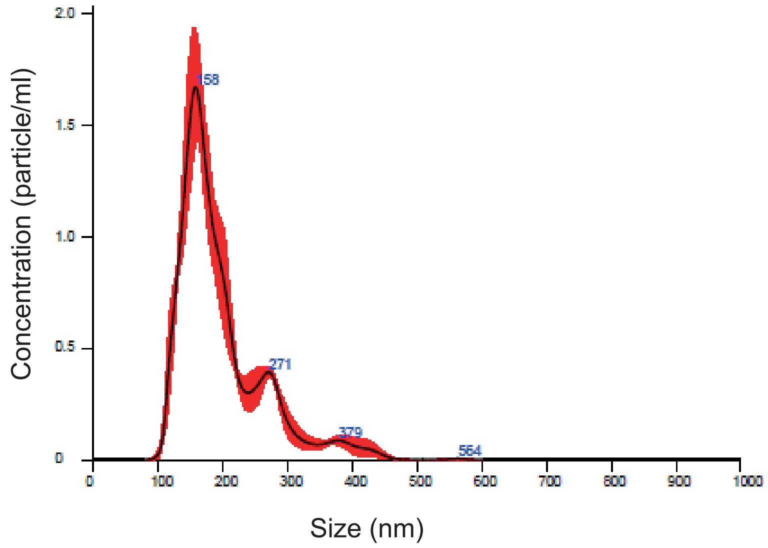

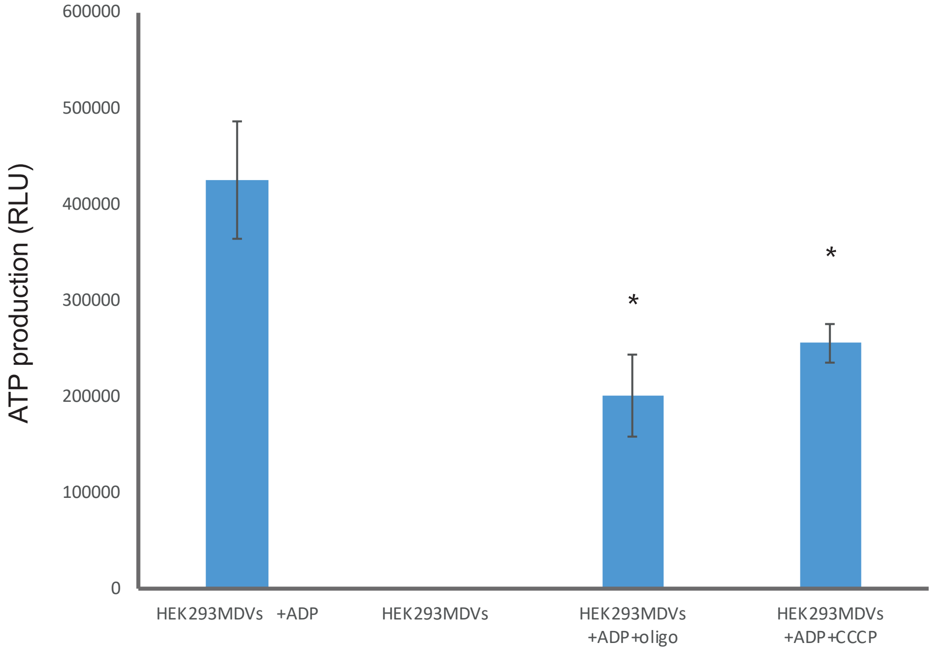

### supplemented figure 3

Supplementary Figure 4

A

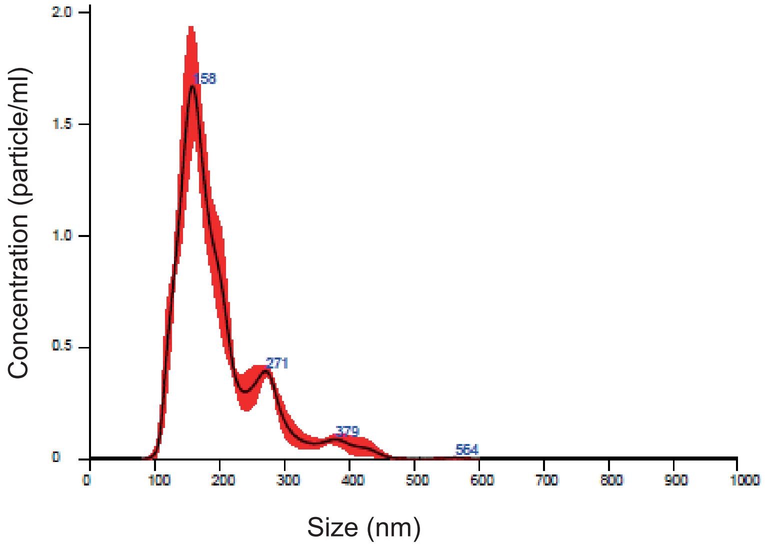


B

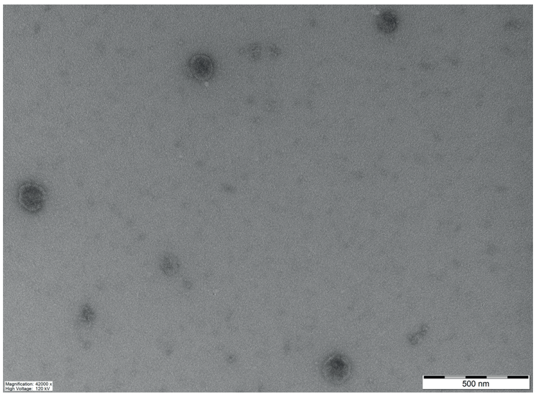


C

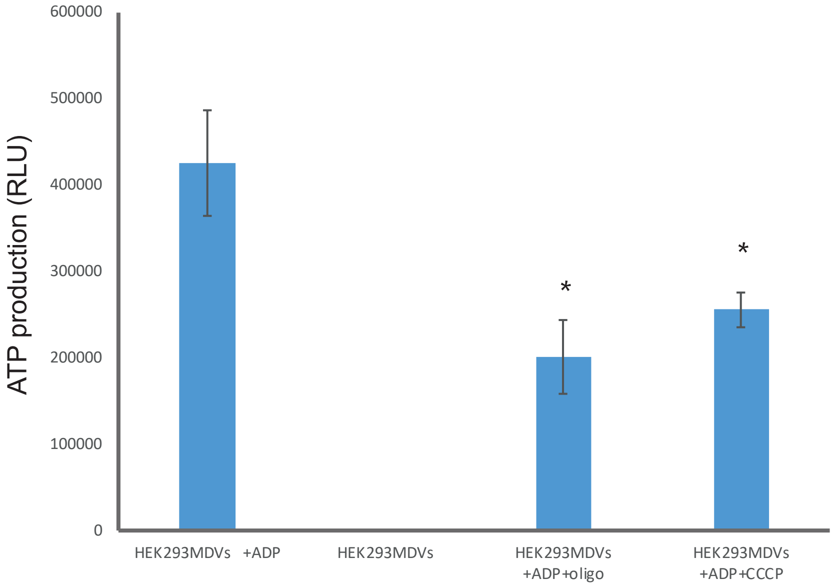
